# Genome-wide mapping of *Vibrio cholerae* VpsT binding identifies a mechanism for c-di-GMP homeostasis

**DOI:** 10.1101/2021.10.04.463056

**Authors:** Thomas Guest, James R.J. Haycocks, Gemma Z.L. Warren, David C. Grainger

## Abstract

Many bacteria use cyclic dimeric guanosine monophosphate (c-di-GMP) to control changes in lifestyle. The molecule, synthesised by proteins having diguanylate cyclase activity, is often a signal to transition from motile to sedentary behaviour. In *Vibrio cholerae*, c-di-GMP can exert its effects via the transcription factors VpsT and VpsR. Together, these proteins activate genes needed for *V. cholerae* to form biofilms. In this work, we have mapped the genome-wide distribution of VpsT in a search for further regulatory roles. We show that VpsT binds 23 loci and recognises a degenerate DNA palindrome having the consensus 5’-W_-5_R_-4_[CG]_-3_Y_-2_W_-1_W_+1_R_+2_[GC]_+3_Y_+4_W_+5_-3’. Most genes targeted by VpsT encode functions related to motility, biofilm formation, or c-di-GMP metabolism. Most notably, VpsT activates expression of the *vpvABC* operon that encodes a diguanylate cyclase. This creates a positive feedback loop needed to maintain intracellular levels of c-di-GMP. Mutation of the key VpsT binding site, upstream of *vpvABC*, severs the loop and c-di-GMP levels fall accordingly. Hence, as well as relaying the c-di-GMP signal, VpsT impacts c-di-GMP homeostasis.

## INTRODUCTION

Cholera is an acute diarrhoeal disease caused by the bacterium *Vibrio cholerae* (1). Globally, the current seventh pandemic is caused by strains of the O1 El Tor biotype that emerged in the Bay of Bengal, later spreading across the globe in three waves (2). Once introduced at a locality, *V. cholerae* strains capable of causing disease outbreaks can remain for several decades (3, 4). In part, this is because *V. cholerae* can spend long periods in aquatic environments (5). In this niche, *V. cholerae* forms biofilms on the chitinous exoskeletons of crustaceans and marine microbiota (5). Hence, these abundant surfaces are exploited as a source of carbon and nitrogen (6, 7). Major components of the *V. cholerae* biofilm matrix include <u>v</u>ibrio <u>p</u>oly<u>s</u>accharide (VPS) and three proteins; RbmA, RbmC and Bap-1 (8). Synthesis of VPS depends on the vps-I and vps-II gene clusters that are transcribed from the *vpsA* and *vpsL* promoters respectively. The nearby *rbm* gene cluster encodes RbmA and RbmB whilst Bap-1 is encoded elsewhere (9). Phase variation between smooth and rugose morphotypes also plays a role (10). Notably, rugose cells form biofilms more readily, present as corrugated colonies on agar, and aggregate in liquid culture (11, 12).

In many bacteria, the decision to form biofilms involves the nucleotide second messenger cyclic dimeric guanosine monophosphate (c-di-GMP) (13). Within cells, c-di-GMP is produced by diguanylate cyclase (DGC) proteins. Such enzymes are identifiable by virtue of a “GGDEF” active site amino acid motif (14). Phosphodiesterases, with “EAL” or “HD-GYP” domains, degrade c-di-GMP (15). The signalling molecule is detected by factors that modulate cell behaviour. For instance, c-di-GMP controlled transcription factors can alter global patterns of gene expression (13). Similarly, the message can be relayed by riboswitches and sRNA molecules (16–18). In *V. cholerae*, two c-di-GMP dependent regulators, VpsR and VpsT, activate the vps-I, vps-II and *rbm* gene clusters (19–21). In rugose phase variants, VpsR and VpsT are required to maintain the phenotype (22, 23). The wider role of these regulators in the direct control of gene expression is less well understood.

First described in 2004, VpsT is a LuxR-type response regulator (20, 22, 24). Direct interaction with c-di-GMP is required for VpsT dimerisation and DNA binding (20). At the *vpsA, vpsL* and *rbm* promoters, VpsT displaces the histone-like nucleoid structuring (H-NS) protein that otherwise silences transcription (25). Conversely, at other loci, the transcription activation mechanism is unknown (26, 27). Transcriptional repression can also be mediated by VpsT. For instance, gene expression is blocked by VpsT binding across the *rpoS* transcription start sites (TSSs) (28). Similarly, VpsT can inhibit transcription of the divergent *VC1303* and *VC1304* genes (29). The DNA site bound by VpsT is called the T-box (30). Efforts to define a consensus T-box have been hampered by the paucity and variability of known VpsT targets. Initial studies revealed recognition of identical sequences by VpsT at the *vpsL* and *vpsA* promoters (25, 30). However, this sequence is not found at other targets (26, 27, 30). In this work we sought a better understanding of VpsT, its DNA regulatory targets, and their interplay with c-di-GMP signalling.

## RESULTS AND DISCUSSION

### Genome-wide DNA binding by VpsT in *Vibrio cholerae*

We used ChIP-seq to map the binding of VpsT across the *V. cholerae* genome. To facilitate this, *vpsT* was cloned in plasmid pAMNF. The resulting DNA construct, pAMNF-*vpsT*, encodes an N-terminal 3xFLAG-*vpsT* fusion that was used to transform *V. cholerae* strain E7946. Importantly, the amount of 3xFLAG-VpsT produced from pAMNF-*vpsT* (Figure S1, lanes 4-6) is similar to the amount of VpsT generated from the native chromosomal locus in both smooth (lanes 1-3) and rugose (lanes 7-9) cells. In subsequent ChIP-seq analysis, anti-FLAG antibodies were used to isolate regions of DNA bound to VpsT. An overview of the data are shown in Figure 1a. Genes are shown as dark blue lines (outer two tracks) and the VpsT binding signal is in cyan (inner track). Strikingly, VpsT does not bind chromosome II. Conversely, we identified 23 binding peaks for VpsT across chromosome I (Table 1). In previous work, VpsT binding at 7 different promoter regions has been shown *in vitro* (25–28, 30). We recovered 5 of these interactions (adjacent to *vpsU, vpsA, vpsL, rbmA* and *rpoS*) in our ChIP-seq analysis (Figure 1b, Table 1). We did not detect VpsT binding upstream of *tag* or *katB* (26, 27). Of the 23 ChIP-seq peaks for VpsT, 11 mapped to genes or operons expressed differently in response to perturbation of *vpsT* (Table 1) (20). Consistent with expectation, most VpsT peaks locate to non-coding DNA upstream of a gene 5’ end (Figure 1c). The function of each gene flanking a binding peak is illustrated in Figure 1d. Many encode proteins implicated in biofilm formation, c-di-GMP metabolism, or motility. We used electrophoretic mobility shift assays (EMSAs) to test direct binding of VpsT with 8 of the ChIP-seq derived targets (Figure S2a). Binding was confirmed in all cases. Conversely, VpsT did not bind the *Escherichia coli lacZ* promoter, which was used as a negative control (Figure S2b).

**Figure 1:**
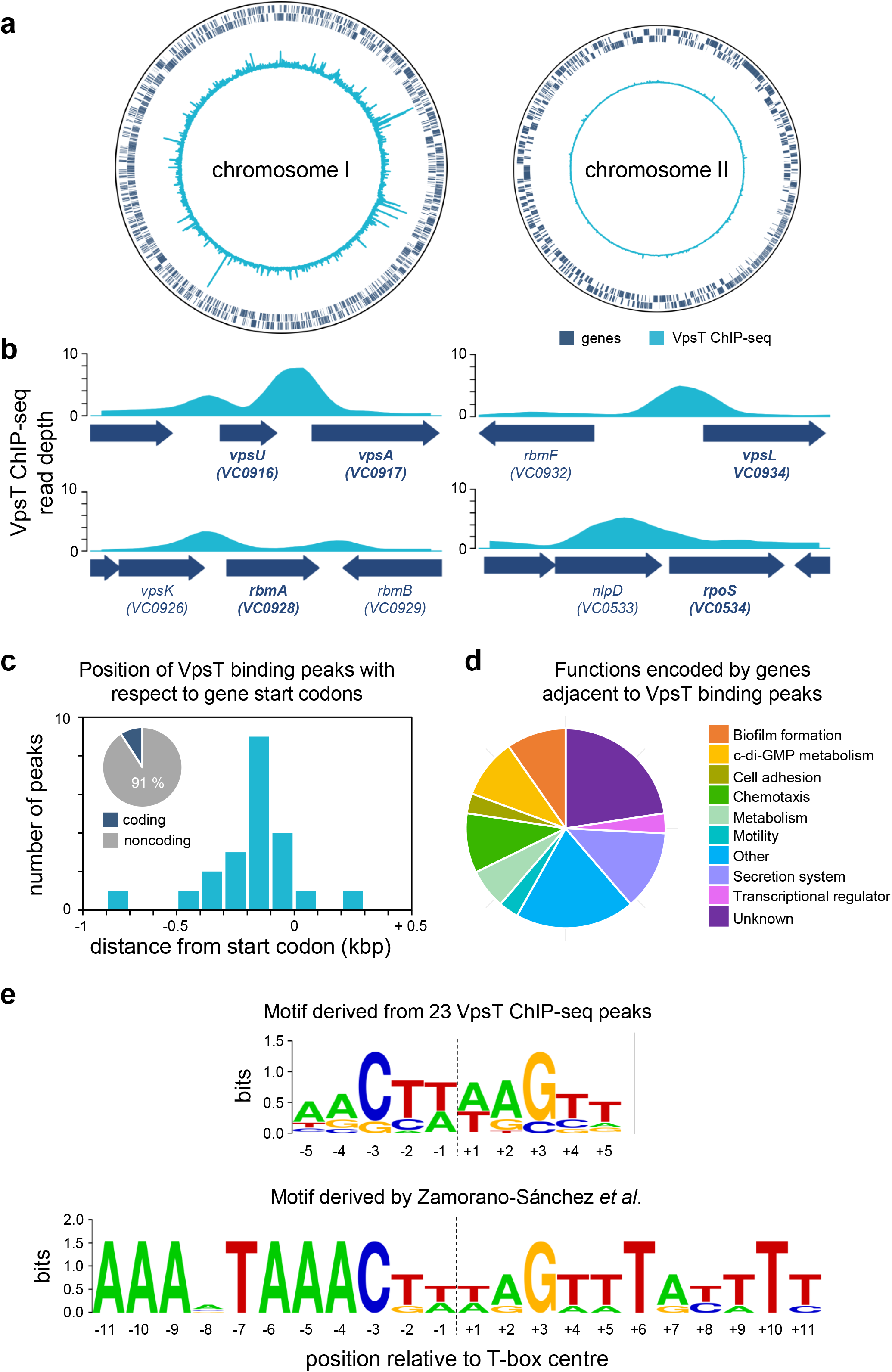
Genome-wide distribution of VpsT in *Vibrio cholerae*. a) Binding of VpsT across both *Vibrio cholerae* chromosomes. The outer two tracks (navy blue) depict genes in the forward and reverse orientation. The VpsT ChIP-seq signal (cyan) is shown by the inner track. b) ChIP-seq coverage plots for known VpsT targets. In each case the VpsT ChIP-seq signal shown in cyan is the average of reads aligned from two independent experiments. Block arrows in navy blue indicate genes that are labelled by name and/or locus tag. Those genes known to be regulated by VpsT are in bold type face. c) Position of VpsT binding peaks with respect to genes. The distribution of VpsT binding peaks occurring at 100 bp intervals from the nearest coding gene. Inset pie chart shows the proportion of peaks that occur in coding and non-coding DNA. d) Pie chart showing the predicted function of genes adjacent to VpsT ChIP-seq peaks. e) DNA sequence logos representing the VpsT binding sequence, or T-box, motif. The upper logo is derived from all 23 VpsT ChIP-seq peaks and the lower sequence was previously reported by Zamorano-Sánchez *et al* (30).

**Table 1:**
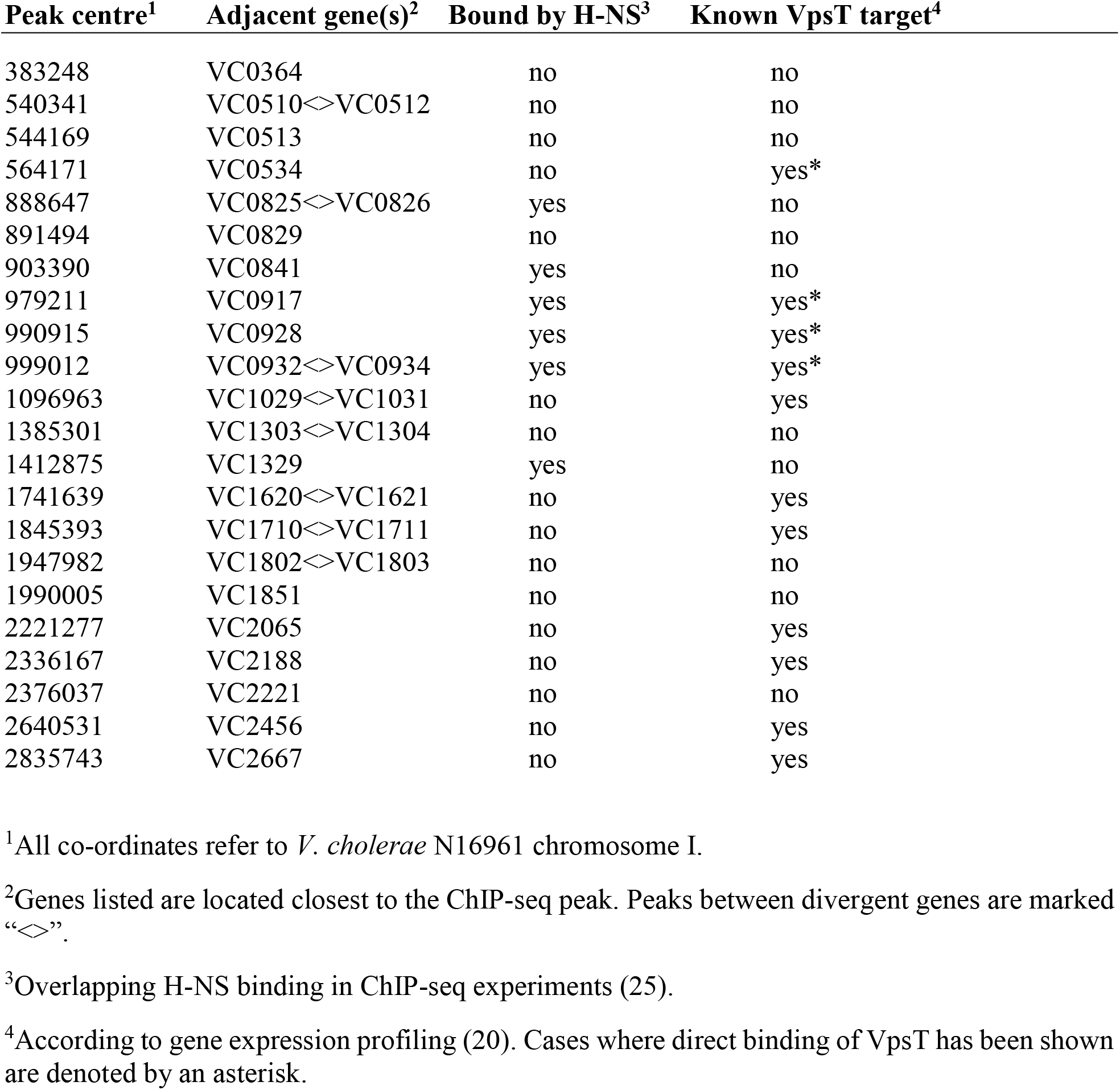
VpsT binding peaks identified by ChIP-seq.

### Reassessment of the T-box consensus sequence for VpsT binding

The precise sequence of the T-box, the DNA motif recognized by VpsT, is not clear. In initial DNAse I footprinting studies, Zamorano-Sánchez *et al*. showed binding of VpsT to a DNA palindrome at the *vpsL* regulatory region (30). A near identical sequence was identified at the *vpsA, vpsT, rbmA* and *rbmE* promoters (30). Direct binding of VpsT has also been demonstrated at the *rpoS* locus using DNAse I footprinting (28). However, the bound sequence differed to the motif previously found (28). Other direct VpsT targets also lack the palindrome identified by Zamorano-Sánchez *et al*. (26, 27, 29). Logically, our ChIP-seq analysis should provide a starting point for better understanding of the T-box consensus. To identify motifs common to all ChIP-seq targets we used MEME to parse DNA sequences centred on each of the 23 VpsT binding peaks. The motif identified is shown in Figure 1e (top) alongside the palindrome previously described by Zamorano-Sánchez *et al*. (bottom) (30). To aid comparison, we numbered positions in both palindromes with respect to the centre of the two sequences (Figure 1e). Whilst the ChIP-seq derived logo is shorter, and exhibits greater degeneracy, there are similarities between the sequences. Most notably, positions −3C and +3G of both motifs are highly conserved and likely to be important for VpsT binding.

### Mutagenesis of the *vpsL* promoter T-box

Recall that Zamorano-Sánchez *et al*. used DNAse I footprinting to locate VpsT binding upstream of *vpsL* (30). Figure 2a illustrates the *vpsL* intergenic region, numbered with respect to the primary TSS (30). The centre of the VpsT ChIP-seq peak is marked by an asterisk. Positions in bold align to the motif found using ChIP-seq. Overlapping sequences matching the original motif of Zamorano-Sánchez and colleagues are underlined (30). To further understand the requirements for VpsT binding we made a series of point mutations. The base changes, numbered with respect to the *vpsL* TSS, are marked red in Figure 2a. To measure the impact of the mutations we used DNAse I footprinting (Figure 2b). As expected, VpsT bound the *vpsL* promoter region poorly in the absence of c-di-GMP (compare lanes 1 and 2). Conversely, in the presence of c-di-GMP, VpsT could elicit a substantial footprint (lane 3). Mutations −236C (lanes 4 and 5) −233T (lanes 6 and 7) and −220G (lanes 12 and 13) target positions in the Zamorano-Sánchez *et al*. logo but beyond the boundaries of the ChIP-seq derived motif. None of these mutations altered VpsT binding. Mutations −228T and −223T target positions conserved in both motifs. Whilst the −228T mutation had no impact (lanes 8 and 9) the −223T mutation abolished VpsT binding (lanes 10 and 11).

**Figure 2:**
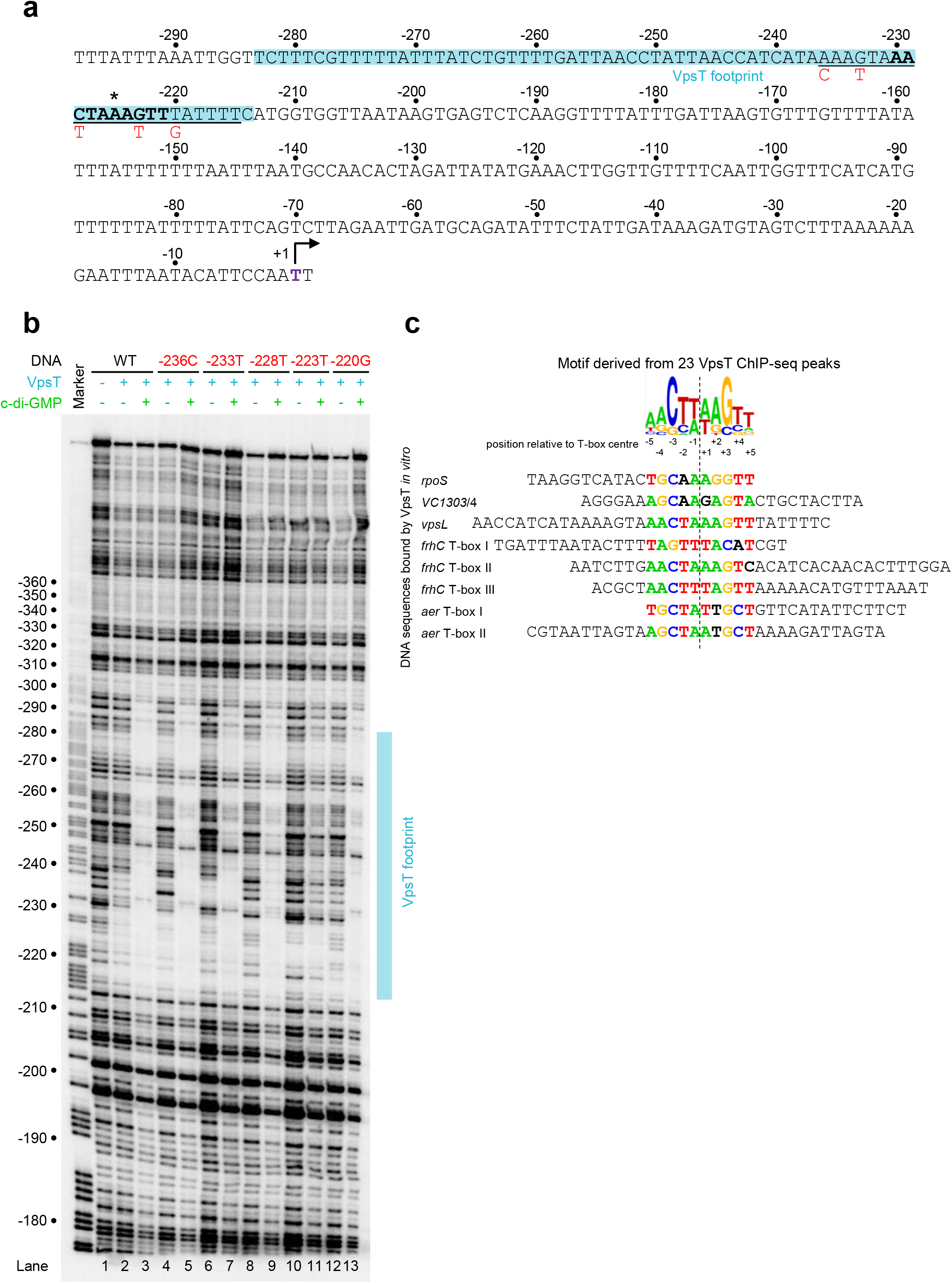
Mutagenesis of the *vpsL* regulatory region T-box. a) DNA sequence of the regulatory region upstream of *vpsL*. The T-box proposed by Zamorano-Sánchez *et al*. is underlined (30). The bold typeface identifies the shorter T-box consensus consistent with all ChIP-seq targets. Mutations made to the T-box are shown below in red. The primary *vpsL* transcription start site (TSS) is indicated by a bent arrow. Regulatory region positions are labelled with respect to the primary *vpsL* TSS (+1). The cyan box highlights the section of the regulatory region protected from DNAse I digestion by VpsT. b) Image of a denaturing polyacrylamide gel used to analyse DNAse I digestion of the *vpsL* regulatory region. The gel is calibrated with a Maxam-Gilbert ‘G+A’ ladder. The presence and absence of VpsT (6 μM) and c-di-GMP (50 μM) is indicated. c) Alignment of DNA sequences protected by VpsT in DNAse I footprinting experiments. Other than for the *rpoS* and *VC1303/VC1304* loci (28, 29), the footprinting data are from the present study. Sequences are aligned according to the position of the T-box (bold type). Aside from *vpsL*, where the sequence was too long, the full protected top strand sequence is shown in the 5’ to 3’ direction. The ChIP-seq derived T-box logo is shown above the aligned sequences for comparison. In the alignment, positions coloured red, green, orange or blue match either the preferred or second most common base according to the ChIP-seq derived motif. Sequences in bold black type match the third or fourth most frequent base.

### Comparison of sequences protected from DNAse I digestion by VpsT

As noted above, prior studies pinpointed VpsT binding, at targets other than *vpsL*, by DNAse I footprinting (28, 29). For comparison, we used the same approach to locate binding of VpsT at the *aer* and *frhC* regulatory regions identified by ChIP-seq (Table 1). At the *aer* locus, VpsT produced two distinct regions of DNA protection that overlapped the centre of the ChIP-seq binding signal (Figure S3). A similar pattern of binding was observed upstream of *frhC*, except that three sections of DNA were bound by VpsT (Figure S4). None of the sequences protected by VpsT, here or in past work, match the T-box sequence identified by Zamorano-Sánchez *et al* (30). We reasoned that the T-box motif identified by ChIP-seq might allow us to resolve these apparent discrepancies. Hence, we collated all eight sequences bound by VpsT in footprinting experiments. We then searched for the ChIP-seq derived T-box motif within these sequences (Figure 3c). In seven cases, an obvious match to the motif was found. In one instance (*frhC* T-box I) a motif was found but the most highly conserved positions, usually −3C and +3G, were inverted (Figure 2c, Figure S4b). Note that the T-box sequence motif suggests that this may occasionally be the case (Figure 1e). We conclude that the ChIP-seq derived logo, generated with the benefit of more VpsT target site information, better describes the DNA binding properties of VpsT.

**Figure 3:**
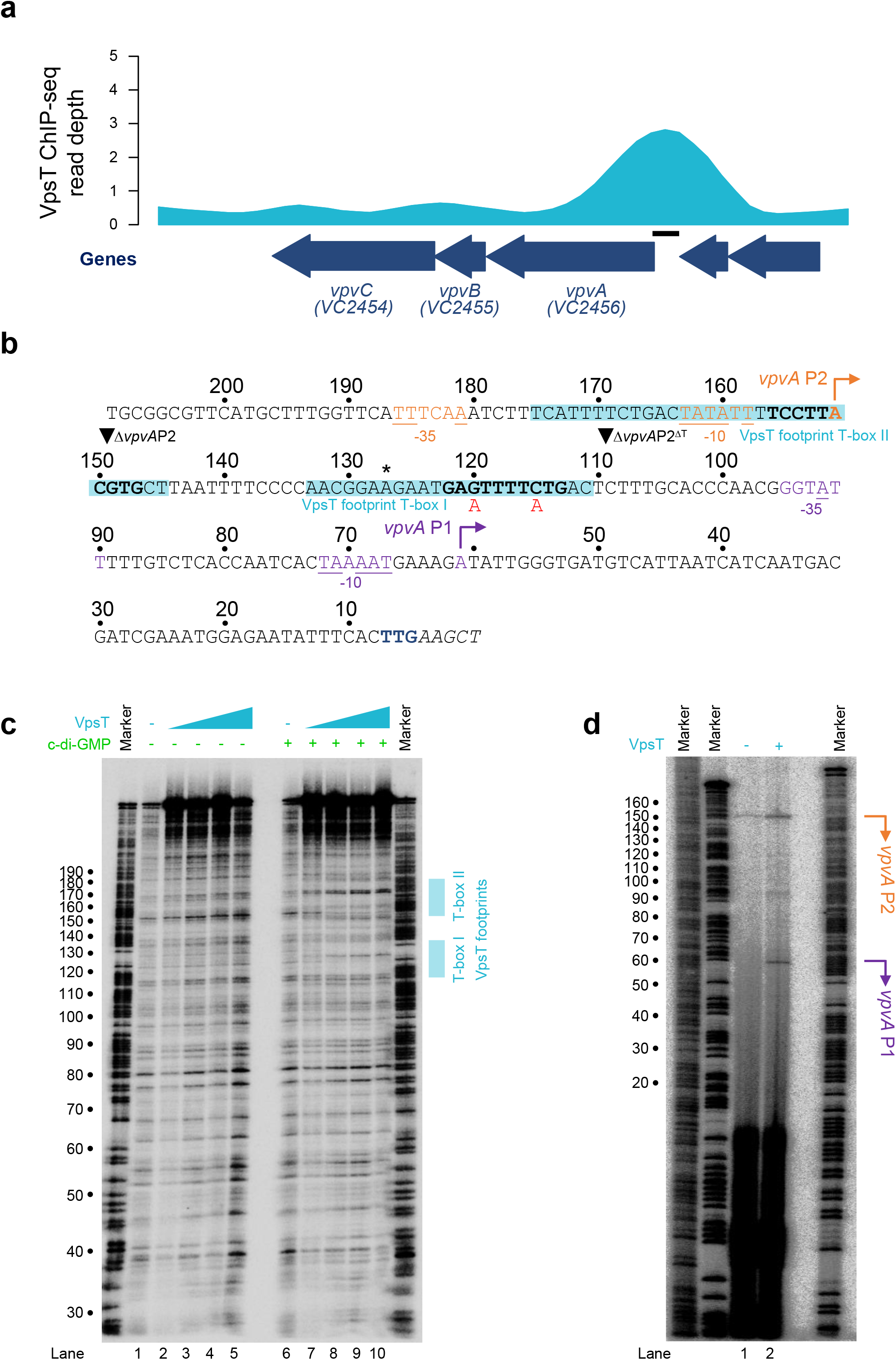
Binding of VpsT upstream of the *vpvABC* operon activates transcription. a) Binding of VpsT to the *vpvABC* regulatory region *in vivo*. The VpsT ChIP-seq signal shown in cyan is the average of reads aligned from two independent experiments. Block arrows in navy blue indicate genes that are labelled by name and/or locus tag. The solid black bar indicates the location of the DNA sequence shown in panel b. b) DNA sequence of regulatory region upstream of *vpvABC*. Bold typeface indicates the T-box consensus identified by ChIP-seq. The asterisk indicates the centre of the ChIP-seq peak for VpsT binding. The cyan box highlights the section of the regulatory region protected from DNAse I digestion by VpsT. Mutations made to the T-box are shown in red. Transcription start sites (+1) are indicated by bent arrows. The −10 and −35 elements of the associated *vpvA*P1 and *vpvA*P2 promoters are also highlighted and matches to the consensus are underlined. Regulatory region positions are labelled with respect to the 3’ end of the DNA fragment used in subsequent DNAse I footprinting experiments. The *vpvA* start codon is coloured navy blue. Inverted triangles indicate sites of truncation to generate derivatives of the regulatory DNA. c) Picture of a denaturing polyacrylamide gel used to separate DNA fragments resulting from DNAse I digestion of the *vpvABC* regulatory region. The gel is calibrated with a Maxam-Gilbert ‘G+A’ ladder. The presence and absence of VpsT (2, 4, 6 or 8 μM) and c-di-GMP (50 μM) is indicated. d) Identification of transcription start sites upstream of *vpv*ABC. The gel image shows the cDNA products of a primer extension assay. Arbitrary Maxam-Gilbert ‘G+A’ ladders were used to calibrate the gel. The RNA used for primer extension was extracted from *V. cholerae* strain E7946 carrying the *vpvABC* regulatory region cloned in plasmid pRW50T. The cells also harboured empty plasmid pAMNF (-VpsT, lane 1) or pAMNF-*vpsT* (+ VpsT, lane 2). Transcription start sites are labelled according to the schematic in panel b.

### VpsT binds two T-boxes upstream of the vibrio phase variation operon

Definition of the T-box consensus should permit easier dissection of gene regulation by VpsT. Hence, we focused our attention on the vibrio phase variation (*vpv*) operon that contains three genes: *vpvA*, *vpvB* and *vpvC*. Importantly, whilst VpvA and VpvB are hypothetical proteins of unknown function, VpvC is a diguanylate cyclase contributing to the corrugated morphology of rugose phase variants (31). The ChIP-seq data for VpsT binding at the *vpvABC* locus is shown in Figure 3a. The sequence of the intergenic region is shown in Figure 3b where the centre of the ChIP-seq peak is indicated by an asterisk. Note that sequences are numbered according to distance from the 3’ end of the DNA sequence. To precisely identify the location of VpsT binding we again used DNAse I footprinting (Figure 3c). In the absence of c-di-GMP no VpsT binding was detected (lanes 1-5). However, in the presence of c-di-GMP, two sections of the regulatory region were protected from DNAse I digestion (lanes 6-10). The regions of protection, each ~20 bp in length, overlap the centre of the ChIP-seq peak and are indicated by cyan boxed sequences in Figure 3b. Examination of the sequences identified T-box motifs (bold in Figure 3b) that we named T-box I and T-box II. Note that T-box I has the same sequence inversion at positions −3 and +3 described above for *frhC* T-box I.

### The regulatory region of the vibrio phase variation operon contains two promoters

To begin understanding the regulatory role of VpsT we searched for TSSs, upstream of *vpvABC*, using primer extension assays. To do this, the DNA sequence shown in Figure 3b was cloned upstream of *lacZ* in plasmid pRW50T. The resulting DNA construct was transferred into *V. cholerae* E7946 by conjugation. These cells were then transformed with pAMNF, or pAMNF-*vpsT*, so that we could determine the consequence of VpsT expression. Finally, RNA was extracted and an oligonucleotide complementary to *lacZ* was used to prime reverse transcription. An electrophoretic analysis of the resulting cDNA products is shown in Figure 3d. In the absence of ectopic VpsT expression, we could detect a single cDNA (Figure 3d, lane 1). This primer extension product indicates a TSS located at position 151 of the cloned *vpvABC* regulatory region (marked orange in Figure 3b). In the presence of VpsT, we could detect an additional cDNA species (Figure 3d, lane 2). The second extension product results from a TSS located at position 61 of the cloned *vpvABC* regulatory region (marked purple in Figure 3b). We named the promoters associated with each TSS *vpvA*P1 and *vpvA*P2, according to their proximity to the *vpvA* start codon (Figure 3b). We note that Papenfort and co-workers previously used dRNA-seq to map TSSs genome-wide in *V. cholerae* (32). This analysis, done using conditions not expected to induce VpsT controlled genes, identified only the *vpvA*P2 TSS. We conclude that *vpv*P1 is a strictly VpsT activated promoter. The constitutive activity of *vpvA*P2 is likely a result of better matches to promoter element consensus sequences (Figure 3b).

### Activation of vpv*A*P1 by VpsT requires the adjacent T-box

To further understand interplay between the promoter and T-box sequences we made derivatives of the *vpvABC* regulatory region. The DNA fragments are illustrated schematically in Figure 4a and the different 5’ ends are marked by inverted triangles in Figure 3b. The various sequences were fused to *lacZ* in plasmid pRW50T and β-galactosidase activity was determined in lysates of *V. cholerae* strain E7946, with or without ectopic VpsT expression. As expected, the starting regulatory fragment was able to stimulate *lacZ* expression and β-galactosidase activity doubled when VpsT was produced (Figure 4b, wild type). To understand the contribution of *vpvA*P2 promoter, and the overlapping T-box II, we removed 60 base pairs from the 5’ end of the wild type DNA fragment. The truncated DNA sequence, named Δ*vpvA*P2, stimulated lower levels of β-galactosidase activity but the fold induction by VpsT was unaltered (Figure 4b, Δ*vpvA*P2). Hence, activation of *vpvA*P1 by VpsT does not require T-box II. To confirm the importance of T-box I we altered the sequence of this DNA element in the context of the truncated DNA fragment. The mutations are indicated by red text in Figure 3b and the resulting DNA fragment was named Δ*vpvA*P2^Mut^. Mutation of T-box I abolished activation of *vpvA*P1by VpsT (Figure 4b, Δ*vpvA*P2^Mut^). In a final experiment, we deleted T-box I completely and activation by VpsT was similarly ablated (Figure 4b, Δ*vpvA*P2^ΔT^).

**Figure 4:**
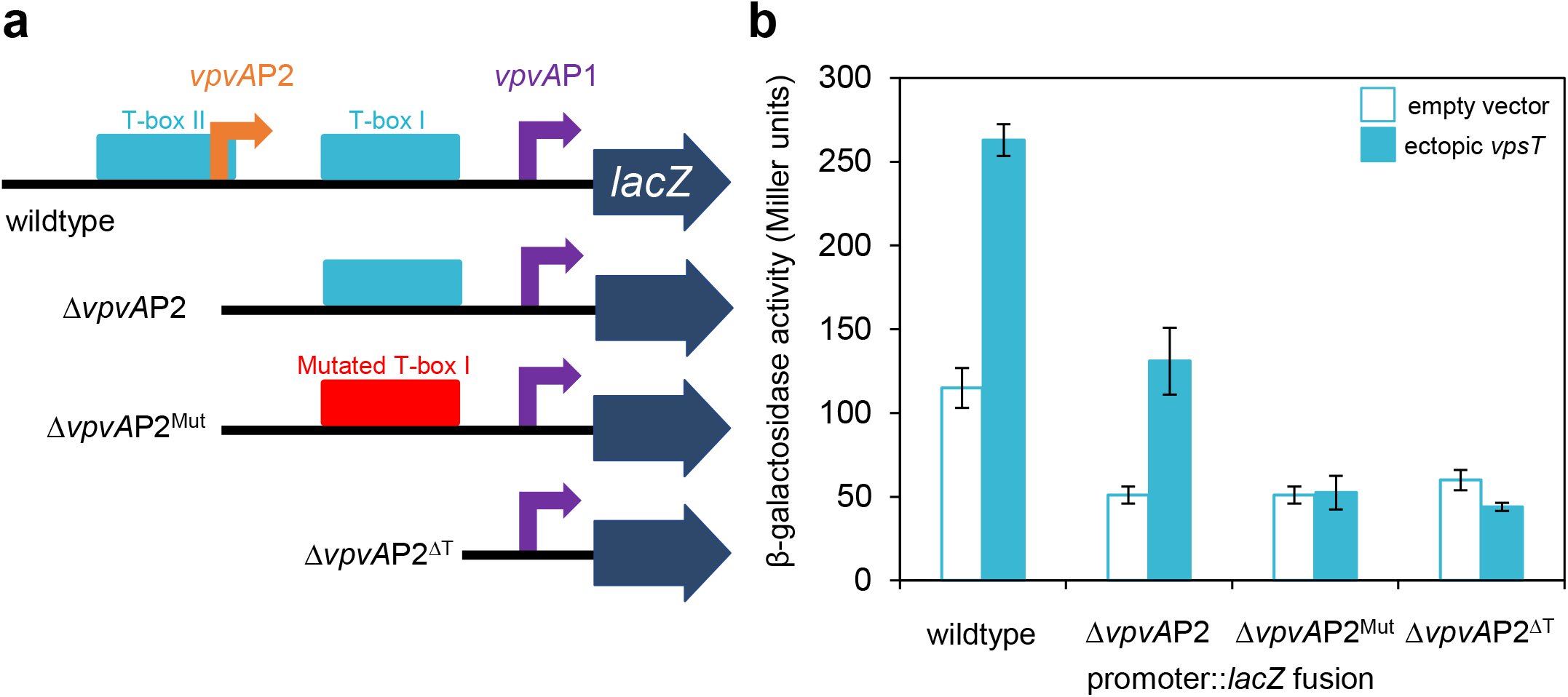
Activation of the *vpvA*P1 promoter requires VpsT and the adjacent T-box. a) Schematic representation of *vpvABC* regulatory region DNA fragments fused to *lacZ*. Truncations are at the sites indicated in Figure 3b and remove T-box II and/or *vpvA*P2 promoter. Transcription start sites are shown by bent arrows and labelled according to the name of the associated promoter. The wild type or mutated T-box I is shown cyan or red respectively. The downstream *lacZ* gene is represented by a block arrow. b) Measurements of β-galactosidase activity in the presence and absence of VpsT. Activities were measured for derivatives of *V. cholerae* strain E7946 encoding the different regulatory region fragments cloned in the *lacZ* reporter plasmid pRW50T. The cells also carried empty plasmid pAMNF (-VpsT) or pAMNF-*vpsT* (+ VpsT). Error bars show standard deviation for three independent biological replicates.

### The *vpvA*P1 promoter and T-box I are needed to maintain intracellular c-di-GMP levels

Previous work has shown that diguanylate cyclase activity generates c-di-GMP (13) and that this molecule positively regulates the activity of VpsT (20). In turn, we show that VpsT activates expression of *vpvABC*, which encodes a diguanylate cyclase. We reasoned that this should constitute a positive feedback loop dependent on T-box I. To test this, we made alterations to the chromosomal *vpvABC* regulatory region. Specifically, we deleted a 60 bp section of chromosomal DNA, containing *vpvA*P2 and T-box II, before making a derivative of this strain with point mutations in T-box I. The chromosomal changes are equivalent to the Δ *vpvA*P2 and Δ*vpvA*P2^Mut^ constructs described above for *lacZ* fusion assays. We also examined the effect of deleting *vpsT* or *vpvABC*. To assess the consequences, we used a plasmid encoded intracellular c-di-GMP biosensor (33). Briefly, the system relies on expression of two different fluorescent reporters. One reporter, AmCyan, is produced constitutively and serves as a normalisation control. The second reporter, TurboRFP, is regulated by a c-di-GMP activated riboswitch. Intracellular c-di-GMP levels are inferred by calculating the AmCyan:TurboRFP ratio. These ratios are shown in Figure 5 alongside images of representative colonies. Wild type colonies have an orange appearance due to the combined expression of AmCyan and TurboRFP. Cells lacking *vpsT* formed colonies with a green-blue appearance due to greatly reduced c-di-GMP levels. Surprisingly, complete deletion of the *vpvABC* operon had no effect. Similarly, deletion of *vpvA*P2 did not induce a large change in c-di-GMP levels. Conversely, mutation of the *vpvABC* T-box I, required for activation of *vpvA*P1 by VpsT, reduced intracellular levels of c-di-GMP by 40 %.

**Figure 5:**
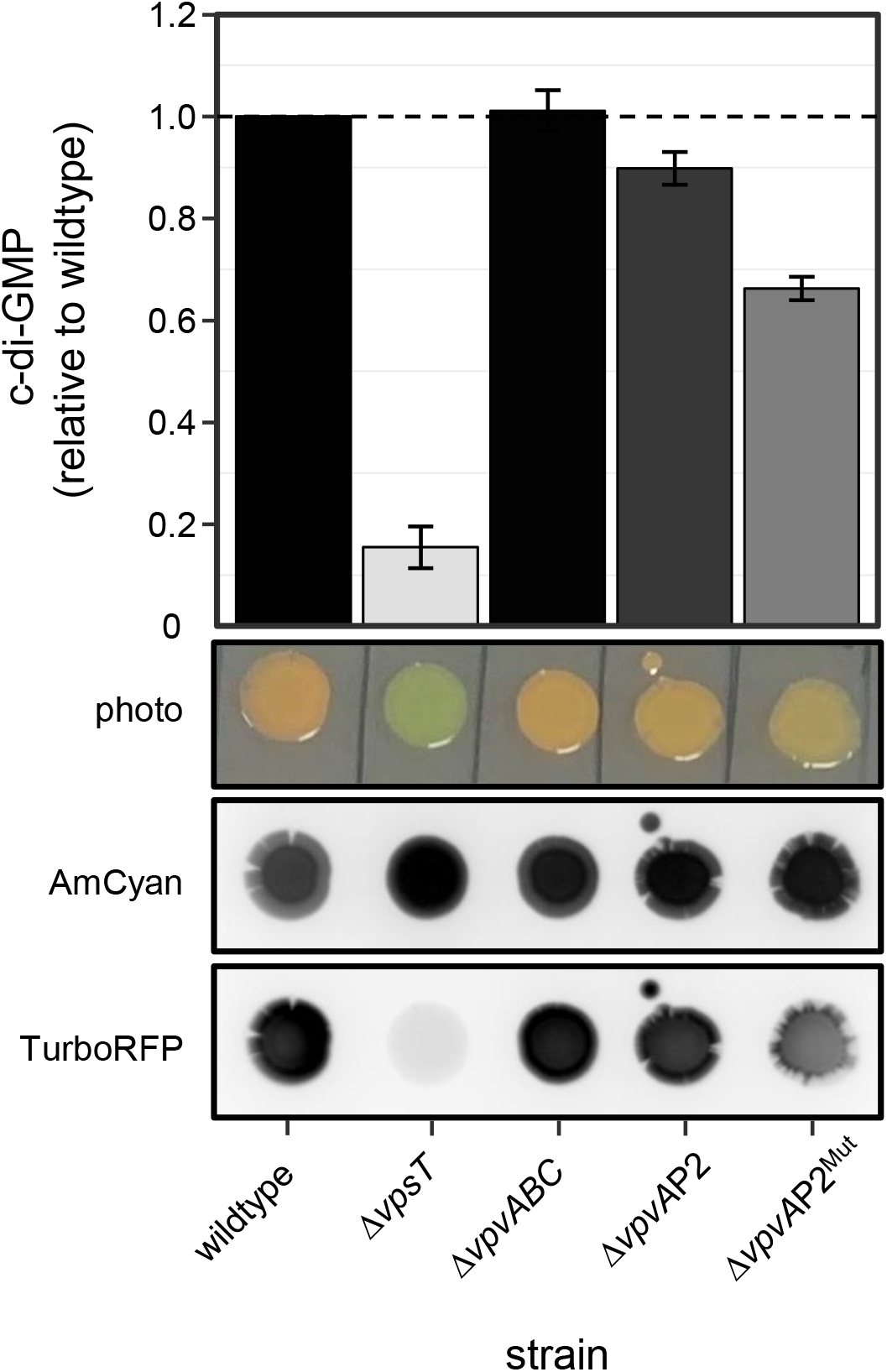
Activation of *vpvABC* by VpsT is needed to maintain intracellular c-di-GMP levels in *Vibrio cholerae*. Relative intracellular concentrations of c-di-GMP were determined in *V. cholerae* strain E7946 and derivatives using a c-di-GMP biosensor (33). The system uses expression of two different fluorescent reporters; constitutive AmCyan expression serves as a normalisation control and TurboRFP expression requires c-di-GMP. Relative intracellular c-di-GMP levels are determined by calculating the TurboRFP:AmCyan ratio for different strains. The two lower panels show representative colonies imaged using blue epi illumination and a 530/28 filter (for AmCyan fluorescence) or green epi illumination and a 605/50 filter (for TurboRFP). A photograph taken in natural light is shown for comparison. The top panel shows a bar chart comparing TurboRFP:AmCyan ratio for different strains. The bars are coloured to match the TurboRFP signal in the bottom image. The strains are distinguished by chromosomal deletions and mutations indicated below the figure. The chromosomal changes at the *vpvABC* regulatory region are equivalent to those described in Figure 4a for promoter:: *lacZ* fusions. Values shown are the average of 4 independent experiments and error bars indicate standard deviation.

**Figure 6:**
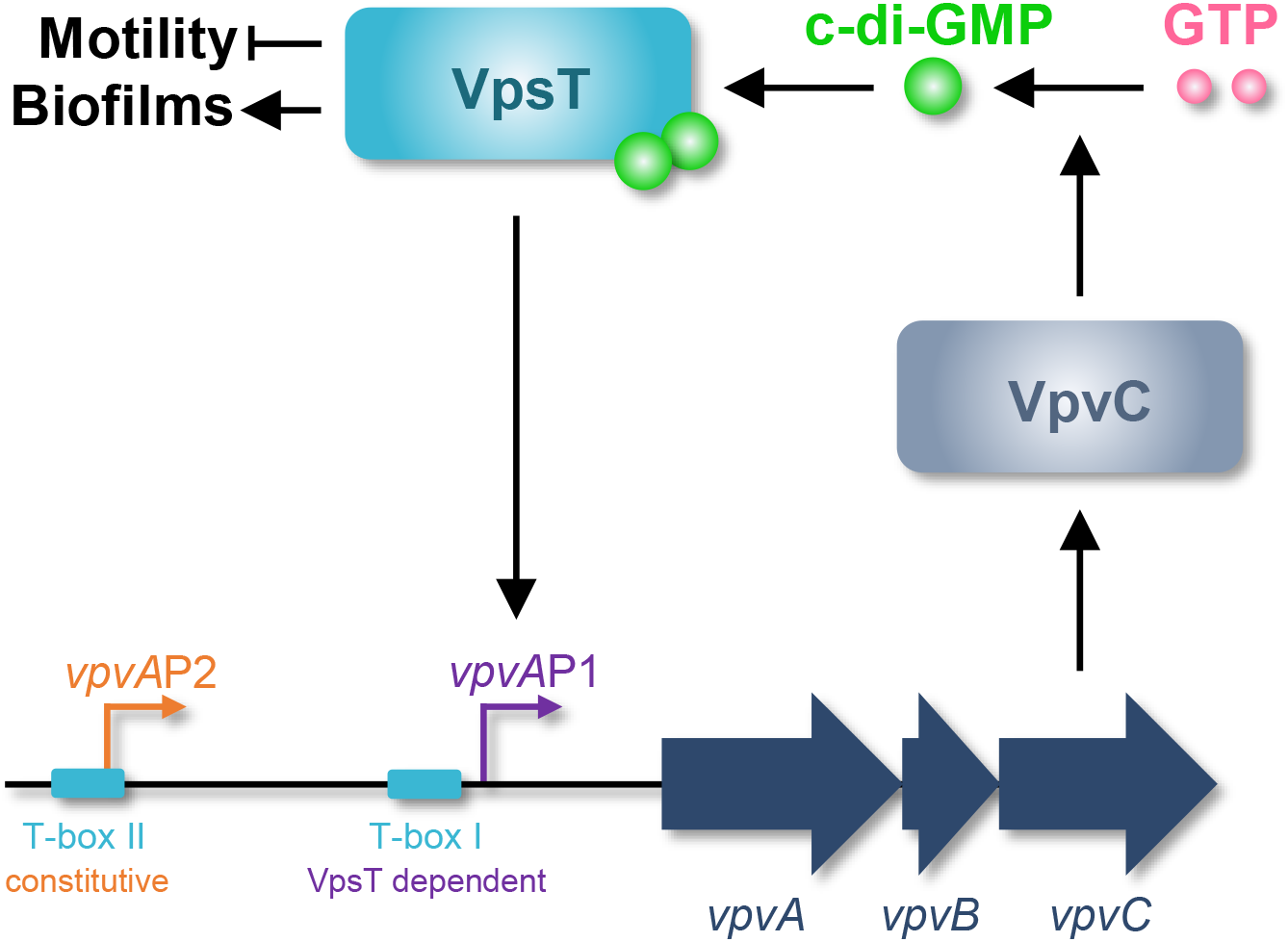
Model describing the different roles of VpsT. Schematic showing the proposed interaction between VpsT, VpvC and c-di-GMP.

### The *vpvA*P2 promoter is important for V. cholerae phase variation

In a final set of experiments, we wanted to understand the contribution of the different *vpvABC* operon regulatory elements to phase variation. Smooth to rugose phase variation in *V. cholerae* can be induced by growing cells in nutrient depleted media (34). We inoculated alkaline phosphate water using a single smooth colony of each strain. After incubation, cells were plated and the number of colonies with the smooth or rugose phenotype were counted. The results are shown in Table 2. As expected, deletion of *vpsT* significantly reduced smooth to rugose phase variation by 4-fold. Conversely, complete deletion of *vpvABC* had no effect. Deletion of *vpvA*P2 caused a significant 2-fold increase in rugose colony numbers that did not change significantly upon deletion of T-box I needed to activate the remaining *vpvA*P1 promoter. We conclude that *vpvA*P2 impacts phase variation but *vpvA*P1 does not.

**Table 2:** Conversion of smooth to rugose variants in different genetic backgrounds.

| **Strain<sup>1</sup>** | **Number of colonies<sup>2</sup>** |  | **% Rugose<sup>3</sup>** | ***p* value<sup>4</sup>** |
| --- | --- | --- | --- | --- |
|  | **Smooth** | **Rugose** |  |  |
| Wild type | 142 | 85 | 37 | N/A |
| $\Delta vpsT$ | 271 | 24 | 8 | <0.00001 |
| $\Delta vpvABC$ | 60 | 42 | 41 | 0.520214 |
| $\Delta vpvAP2$ | 28 | 57 | 67 | <0.00001 |
| $\Delta vpvAP2^{Mut}$ | 36 | 45 | 56 | 0.127989 |

<sup>1</sup>*V. cholerae* strain E7946 or derivatives thereof.
<sup>2</sup>A single smooth colony was used to inoculate alkaline phosphate water. After 72 hours incubation, cells were plated and the number of colonies with the smooth or rugose phenotype were counted.
<sup>3</sup>The percentage of all colonies counted that had a rugose phenotype.
<sup>4</sup>All *p* values were calculated using the Chi-square test. We compared the number of smooth and rugose colonies for the wild type parent strain with each derivative. The exception was the $\Delta vpvAP2^{Mut}$ strain that was compared with $\Delta vpvAP2$ . The null hypothesis was that no differences between strains would be observed.

## CONCLUSIONS

We argue that the T-box is best described as a degenerate palindrome with the consensus 5’-W_-5_R_-4_[CG]_-3_Y_-2_W_-1_W_+1_R_+2_[GC]_+3_Y_+4_W_+5_-3’. This resembles part of a prior T-box motif, generated from a smaller number of targets, and explains observations of VpsT binding diverse sequences (Figure 2c) (28–30). Positions −3 and +3 of the motif are key and, whilst usually C and G respectively, exhibit the complimentary sequence in some cases (Figures 2c and 3b). In both scenarios, mutation of the key positions prevents interaction with VpsT (Figures 2b and 4b). Globally, the DNA binding properties of VpsT are intriguing. Despite recognising a highly degenerate sequence, VpsT targets just 23 loci in our ChIP-seq assay (Table 1). Furthermore, whilst *vpsT* resides within *V. cholerae* chromosome II, all VpsT targets locate to chromosome I. This is largely consistent with prior transcriptome analysis, which identified a similar skew in VpsT regulated genes (20). The unusual distribution is unlikely to be driven solely by occurrence of the T-box sequence; incidence of the motif can be identified frequently across both chromosomes (35). Hence, the ability of VpsT to bind DNA must be influenced by additional factors. Other regulators and DNA supercoiling could both play a role. We note that VpsT appears to bind DNA differently at certain targets. For instance, at the *vpsL* regulatory region, DNA protection stretched ~60 base pairs upstream of the T-box (Figure 2). This protection cannot be due to VpsT independently binding multiple sites; a single T-box point mutation was sufficient to abolish the footprint. Hence, we suggest that VpsT must cooperatively oligomerise across, or tightly wrap, the *vpsL* regulatory DNA. The VpsT binding pattern is different upstream of *vpvABC*; a precise footprint of each T-box is observed. Interestingly, VpsT activates *vpsL* transcription by antagonising H-NS mediated repression (25, 30). This is not the case for *vpvABC*, that does not bind H-NS (25). Hence, we speculate that the more extensive interactions of VpsT at the *vpsL* locus may be necessary to displace H-NS.

Our data suggest that the *vpvA*P1 and *vpvA*P2 promoters have different roles. The latter is constitutively active whilst *vpvA*P1 is activated by VpsT (Figure 3) (32). The constitutive activity of *vpvA*P2 is consistent this promoter impacting the rate of phase variation; rugose colonies can result from mutation of *vpvC*, basal expression of which requires *vpvA*P2 (Figure 3d) (31, 32). Conversely, the VpsT activated *vpvA*P1 promoter did not impact phase variation (Table 2). However, this promoter did impinge on c-di-GMP homeostasis. This is likely a result of a positive feedback loop; c-di-GMP activates VpsT that subsequently upregulates expression of VpvC and so diguanylate cyclase activity. Hence, mutation of T-box I, needed to activate *vpvA*P1, led to a 40 % decrease in intracellular c-di-GMP (Figure 5). Deletion of *vpsT* caused the intracellular c-di-GMP signal to decrease dramatically (Figure 5). However, this phenotype may depend on regulatory targets in addition to *vpvABC*. The stability of the intracellular c-di-GMP signal following *vpvABC* deletion suggests other DGC proteins play a compensatory role. Indeed, *V. cholerae* encode thirty one proteins with the “GGDEF” motif and redundancy is common (15, 36). Furthermore, as the roles of *vpvA* and *vpvB* are less well defined, their deletion could have confounding consequences. In conclusion, VpsT both relays the c-di-GMP signal and is important for the control of intracellular c-di-GMP levels.

## MATERIALS AND METHODS

### Strains, plasmids and oligonucleotides

Strains, plasmids and oligonucleotides used in this study are listed in Table S1. All *V. cholerae* strains are derivatives of E7946 (37). Chromosomal deletions and mutants were made using either the pKAS32 suicide plasmid for allelic exchange or the MuGENT method (38, 39). The *Escherichia coli* strain JCB387 was used for routine cloning (40). Plasmids were transferred into *V. cholerae* by either conjugation or transformation as described previously (41, 42).

### ChIP-seq and bioinformatics

The chromatin immunoprecipitation assay was done as in prior work (42) using strain E7946, carrying plasmid pAMNF-*vpsT*, or empty pAMNF as a control. Note that pAMNF-*vpsT* encodes 3xFLAG-VpsT that is expressed from a low level constitutively active promoter (43). The VpsT-DNA complexes were immunoprecipitated with an anti-FLAG antibody (Sigma) and Protein A sepharose beads. Immunoprecipitated DNA was blunt-ended, A- tailed, and ligated to barcoded adaptors before elution and de-crosslinking. ChIP-seq libraries were then amplified by PCR and purified. Library quality was assessed using an Agilent Tapestation 4200 instrument and quantity determined by qPCR using an NEBnext library quantification kit (NEB). Libraries were sequenced as described previously (43) and reads are available from ArrayExpress using accession code E-MTAB-10829. Single-end reads from two independent VpsT ChIP-seq experiments were mapped to the reference *V. cholerae* N16961 genome (chromosome I: NC_002505.1 and chromosome II: NC_002506.1) with Bowtie 2 using the QuasR package. MACS2 was used to call ChIP-seq peaks, which were then visually inspected using the Artemis genome browser to confirm peak centres (44, 45). Average read counts were normalised according to the coverage per base for each experiment and visualised using Gviz.

### β-galactosidase activity assays

Promoter DNA was fused to *lacZ* in plasmid pRW50T that can be transferred from *E. coli* to *V. cholerae* by conjugation (41). Assays of β-galactosidase activity were done according to the Miller method (46). Bacterial cultures were grown in LB broth, supplemented with appropriate antibiotics, to mid-log phase. Values shown are the mean of three independent experiments and error bars show the standard deviation.

### Electrophoretic mobility shift assays and DNAse I footprinting

We purified VpsT as described previously (29). Promoter DNA fragments were excised from plasmid pSR and end-labelled with γ32-ATP using T4 PNK (NEB). EMSAs and DNase I footprints were done as previously described (42). Full gel images are shown in Figure S5.

### Primer extension assays

The RNA for primer extension assays was purified from *V. cholerae* carrying appropriate plasmids as described above. Primer extension assays were done as previously described (47). The 5’ end-labelled primer D49724, which anneals downstream of the *Hind*III site in pRW50T, was used in all experiments. Primer extension products were analysed on denaturing 6% polyacrylamide gels, calibrated with arbitrary size standards, and visualized using a Fuji phosphor screen and Bio-Rad Molecular Imager FX. Full gel images are shown in Figure S5.

### Colony fluorescence quantification

Cultures of *V. cholerae* were grown aerobically overnight in LB. A 2 μL suspension was spotted onto LB agar plates that were incubated overnight at 30 °C. The fluorescence signal from each of the two fluorophores was measured with a ChemiDoc MP system (Bio-Rad) as previously described (33). Image Lab (BioRad) was used to measure the fluorescence intensity, with the same number of pixels recorded for each colony. The ratio of TurboRFP to the AmCyan signal was then calculated. Values shown are mean ratios from 4 independent experiments, as a proportion of the wild type ratio.

## ACKNOWLEDGEMENTS

This work was funded by BBSRC project grant BB/N005961/1 awarded to DCG and a BBSRC MIBTP studentship (BB/M01116X/1, project reference 1790836) awarded to TG. We thank Fitnat Yildiz for the gift of plasmid pFY4535 and critical reading of the manuscript prior to submission.

## SUPPLEMENTARY FIGURE LEGENDS

**Figure S1:**
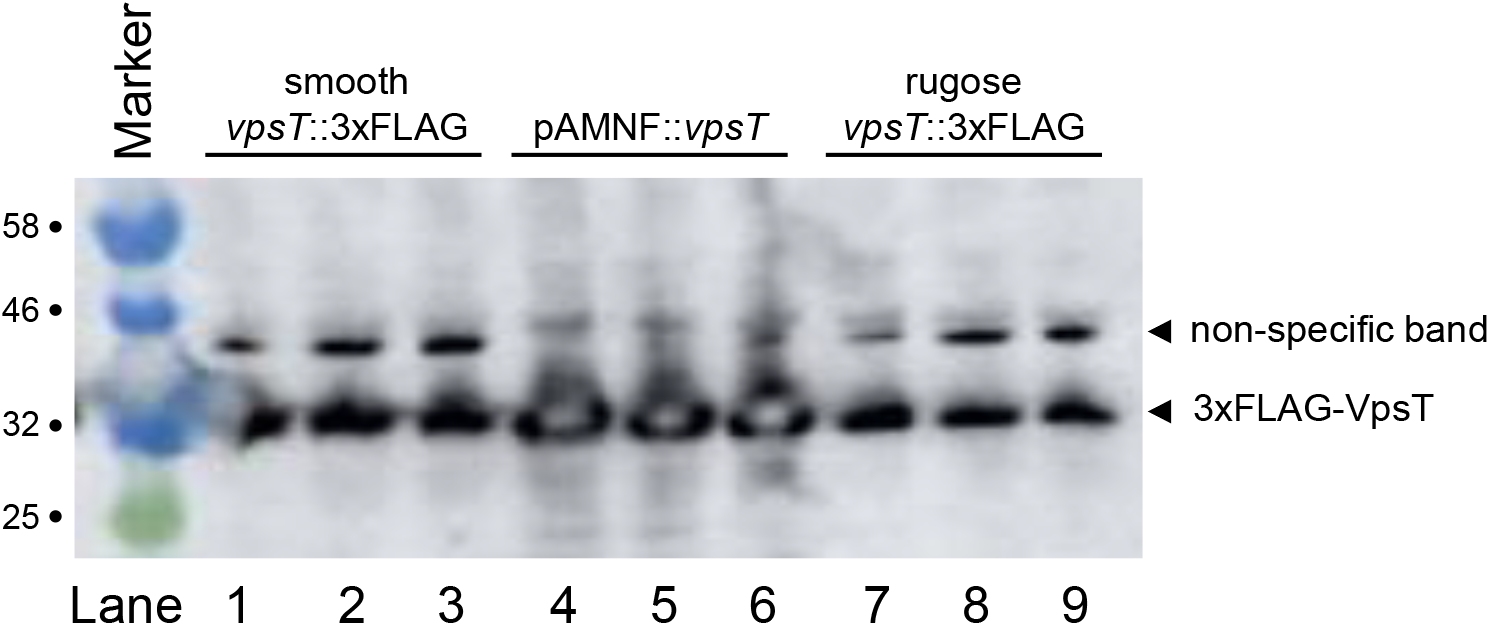
Comparison of VpsT levels in different strains. The figure shows the result of a western blotting experiment, with anti-FLAG antibodies, to detect chromosomal or plasmid encoded 3xFLAG-VpsT. Each measurement was done in triplicate for the strains indicated above the gel image.

**Figure S2:**
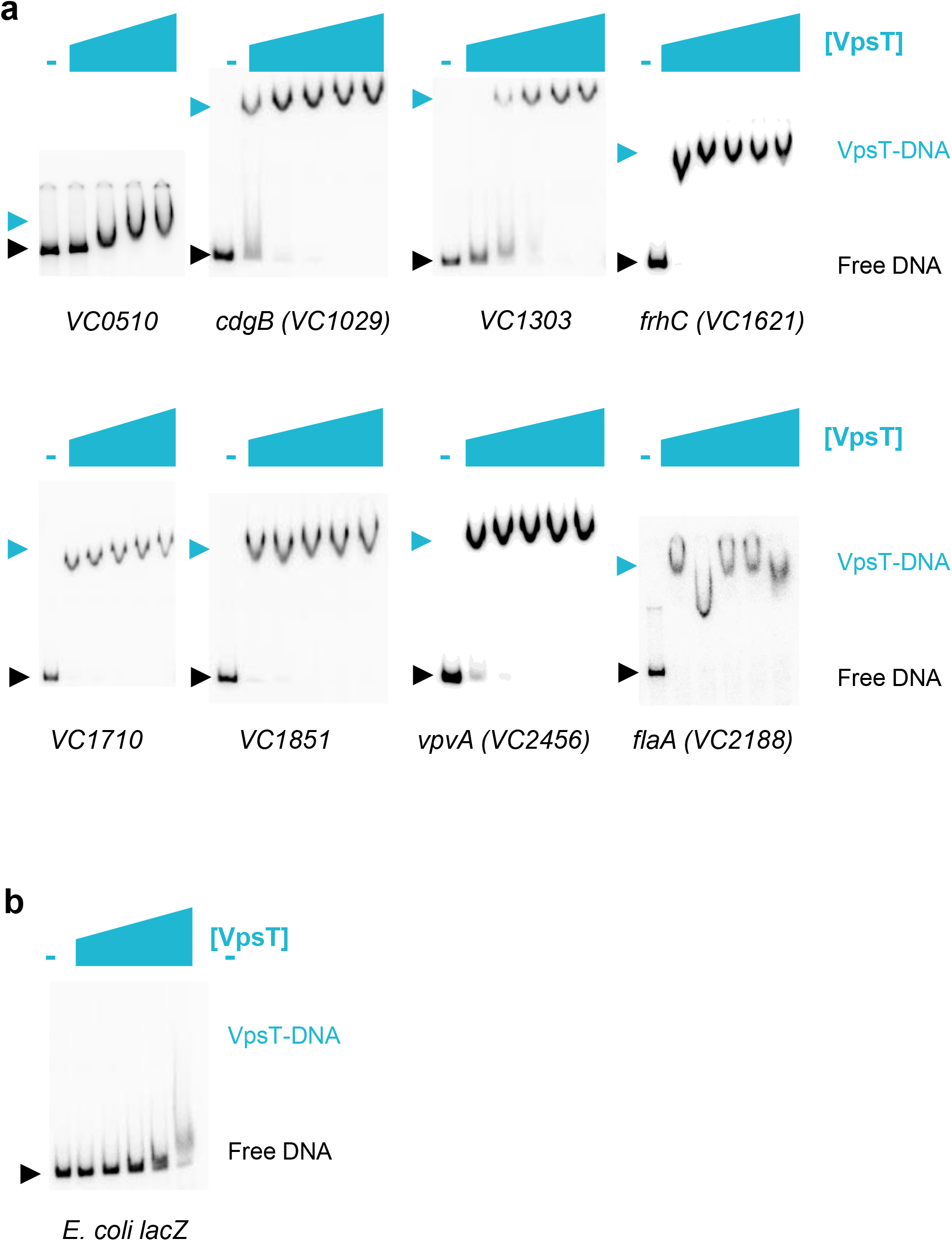
Binding of purified VpsT *in vitro* to DNA targets identified by ChIP-seq *in vivo*. a) The images show results from electrophoretic mobility shift assays with purified VpsT (0-7.5 μM), c-di-GMP (50 μM) and DNA fragments corresponding to ChIP-seq peaks detected for VpsT binding *in vivo*. b) A negative control electrophoretic mobility shift assay demonstrating no binding of VpsT to *Escherichia coli lacZ* DNA.

**Figure S3:**
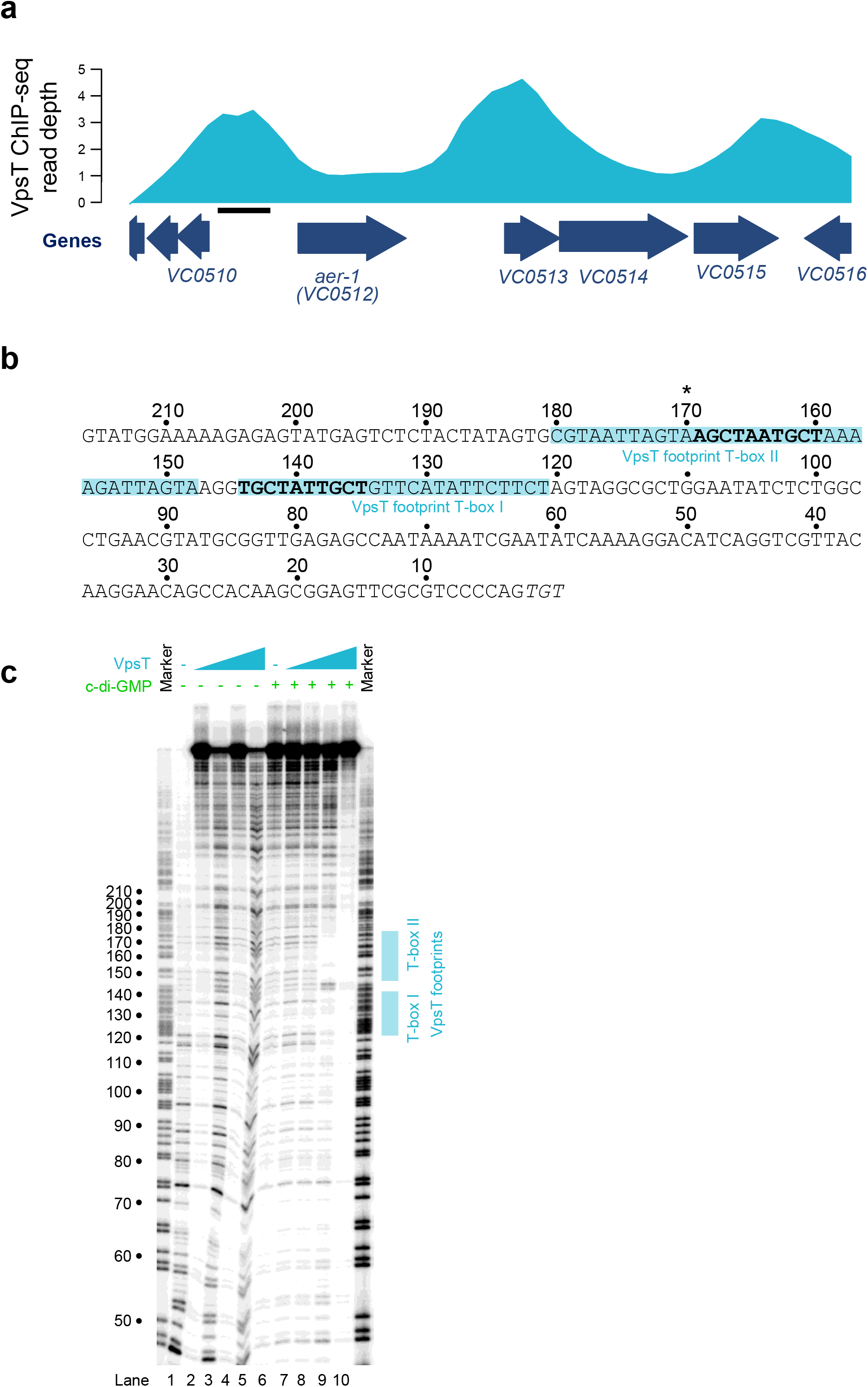
VpsT binds two targets at the *aer* regulatory region. a) Binding of VpsT to the *aer* regulatory region *in vivo*. The VpsT ChIP-seq signal shown in cyan is the average of reads aligned from two independent experiments. Block arrows in navy blue indicate genes that are labelled by name and/or locus tag. The solid black bar indicates the location of the DNA sequence shown in panel b. b) DNA sequence of regulatory region upstream of *aer*. Bold typeface indicates the T-box consensus identified by ChIP-seq. The asterisk indicates the centre of the ChIP-seq peak for VpsT binding. The cyan box highlights the section of the regulatory region protected from DNAse I digestion by VpsT. c) Image of a denaturing polyacrylamide gel used to separate DNA fragments resulting from DNAse I digestion of the *aer* regulatory region. The gel is calibrated with a Maxam-Gilbert ‘G+A’ ladder. The presence and absence of VpsT (2, 4, 6 or 8 μM) and c-di-GMP (50 μM) is indicated.

**Figure S4:**
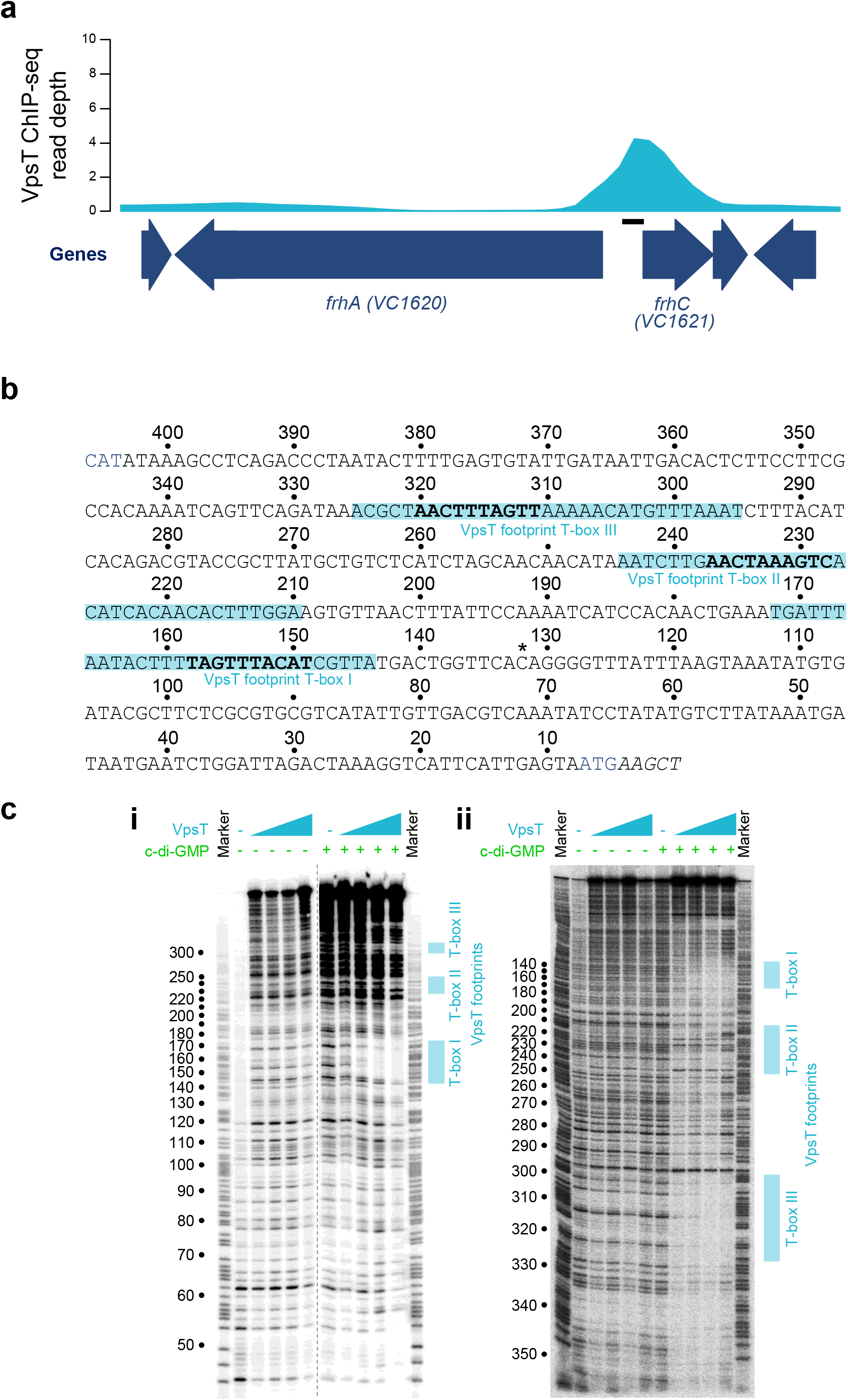
VpsT binds three targets at the *frhA/frhC* regulatory region. a) Binding of VpsT to the *frhA*/*frhC* regulatory region *in vivo*. The VpsT ChIP-seq signal shown in cyan is the average of reads aligned from two independent experiments. Block arrows in navy blue indicate genes that are labelled by name and/or locus tag. The solid black bar indicates the location of the DNA sequence shown in panel b. b) DNA sequence of regulatory region upstream of *frhC*. Bold typeface indicates the T-box consensus identified by ChIP-seq. The asterisk indicates the centre of the ChIP-seq peak for VpsT binding. The cyan box highlights the section of the regulatory region protected from DNAse I digestion by VpsT. The *frhC* start codon is in blue text. c) Image of a two denaturing polyacrylamide gels used to separate DNA fragments resulting from DNAse I digestion of the DNA sequence upstream of *frhC*. The length of the regulatory DNA meant that it was necessary to label the 5’ ends of either the (i) bottom or (ii) top DNA strands to get full coverage of the sequence. The gel is calibrated with a Maxam-Gilbert ‘G+A’ ladder. The presence and absence of VpsT (2, 4, 6 or 8 μM) and c-di-GMP (50 μM) is indicated. c) Image of a denaturing polyacrylamide gel used to separate DNA fragments resulting from DNAse I digestion of the *aer* regulatory region. The gel is calibrated with a Maxam-Gilbert ‘G+A’ ladder. The presence and absence of VpsT (2, 4, 6 or 8 μM) and c-di-GMP (50 μM) is indicated.

**Figure S5:**
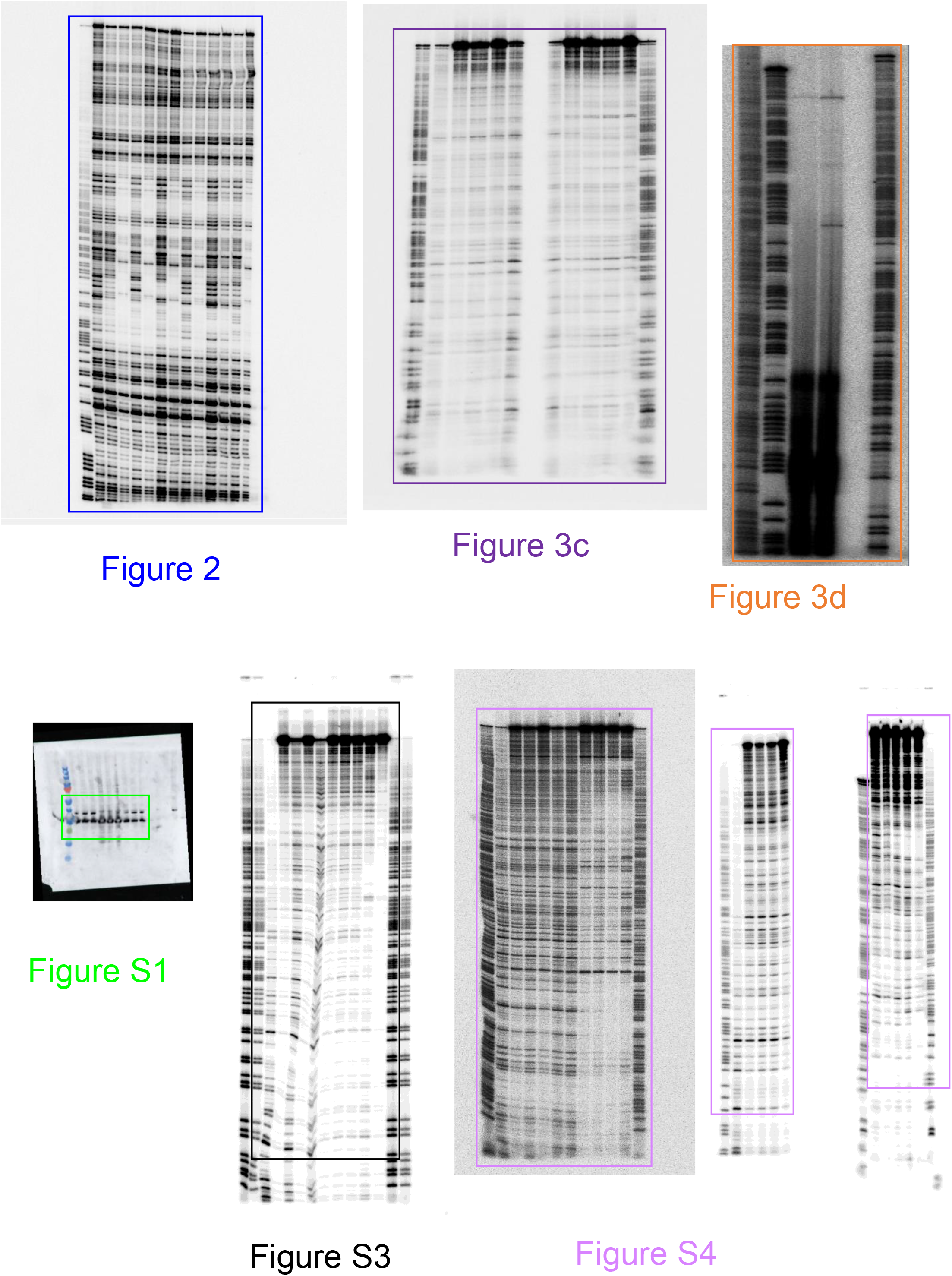
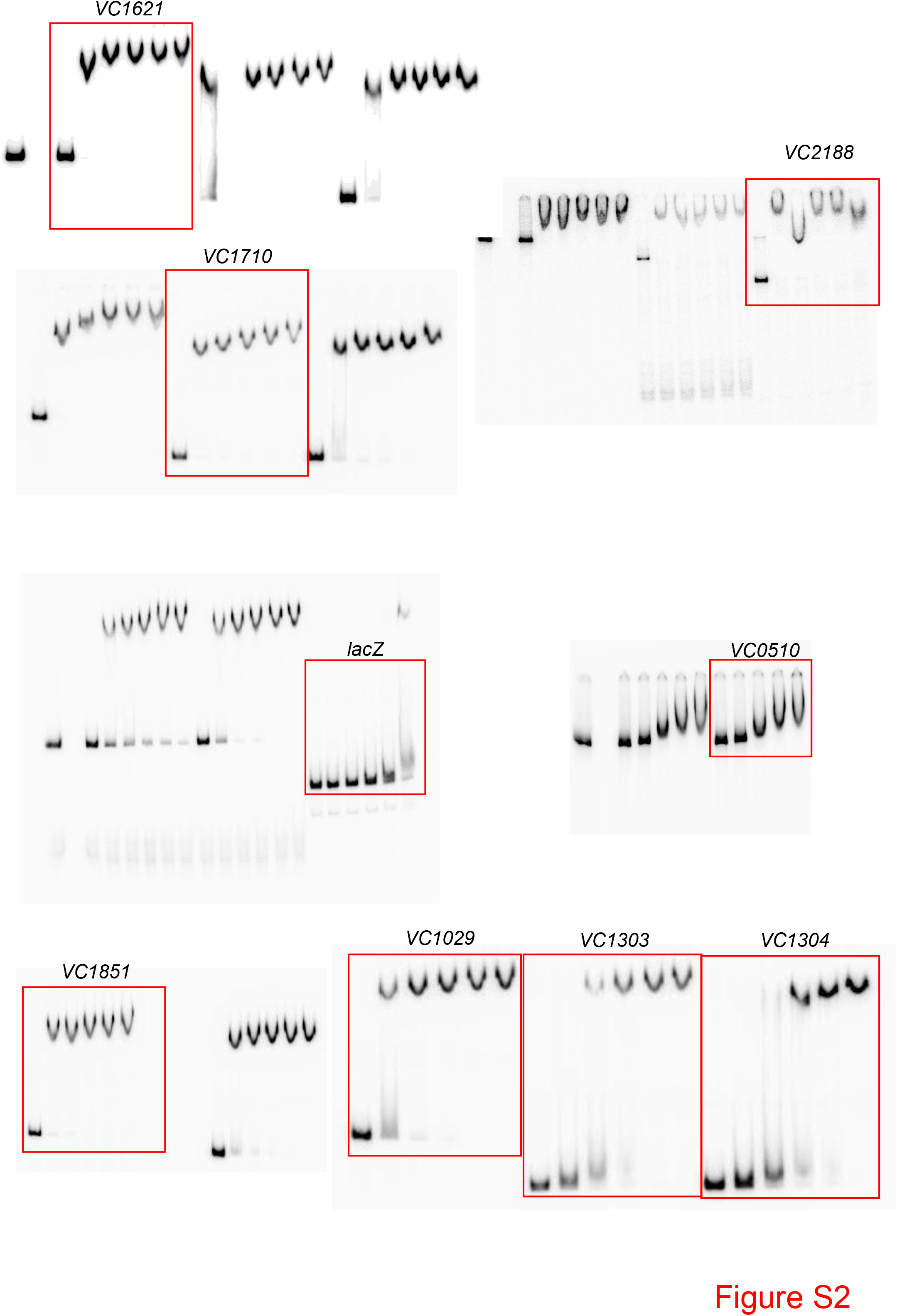
Original gel images. Original uncropped gel images used for this work. Coloured boxes indicate the approximate boundaries of images following cropping of images for figures.

---

**Table S1:**
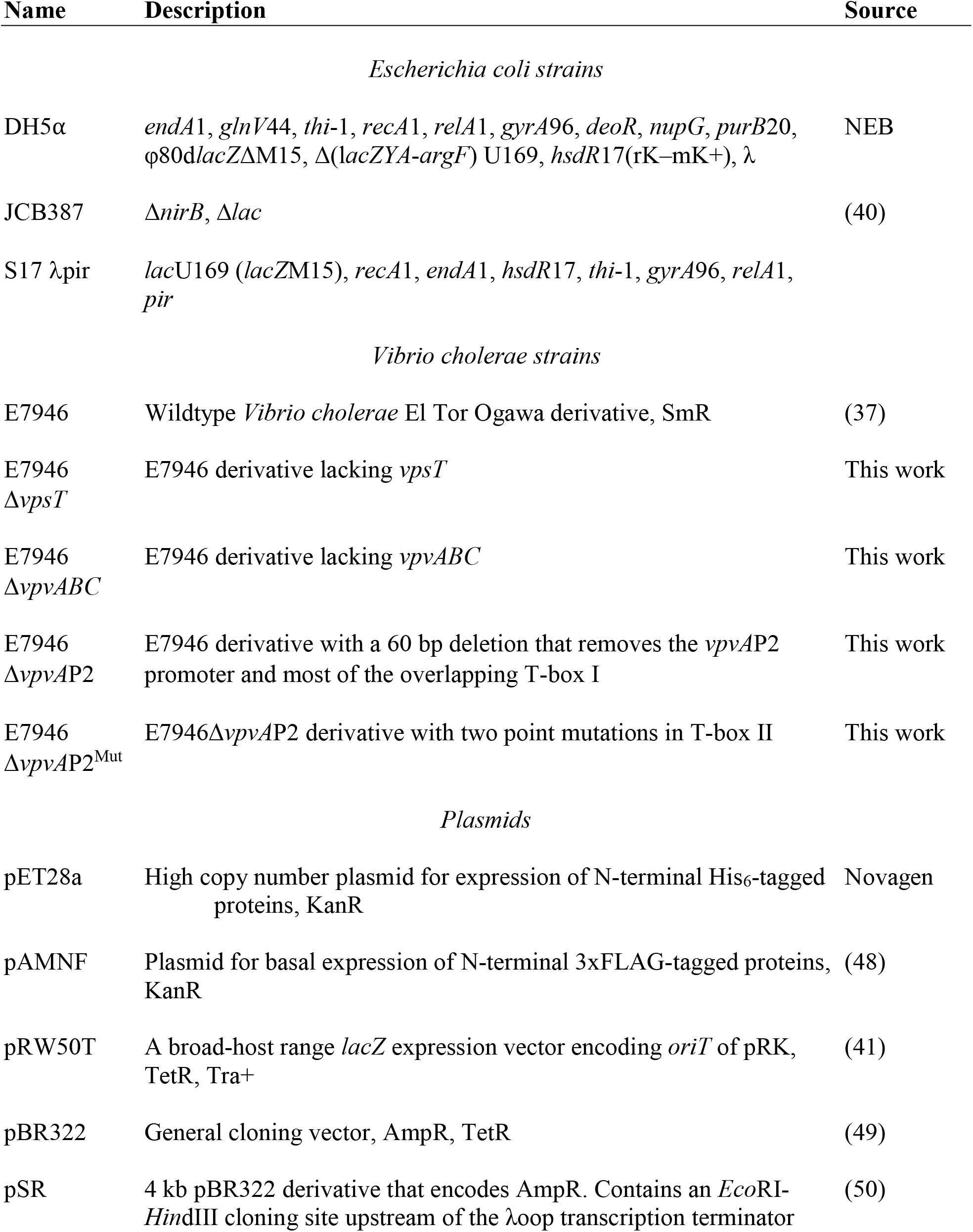
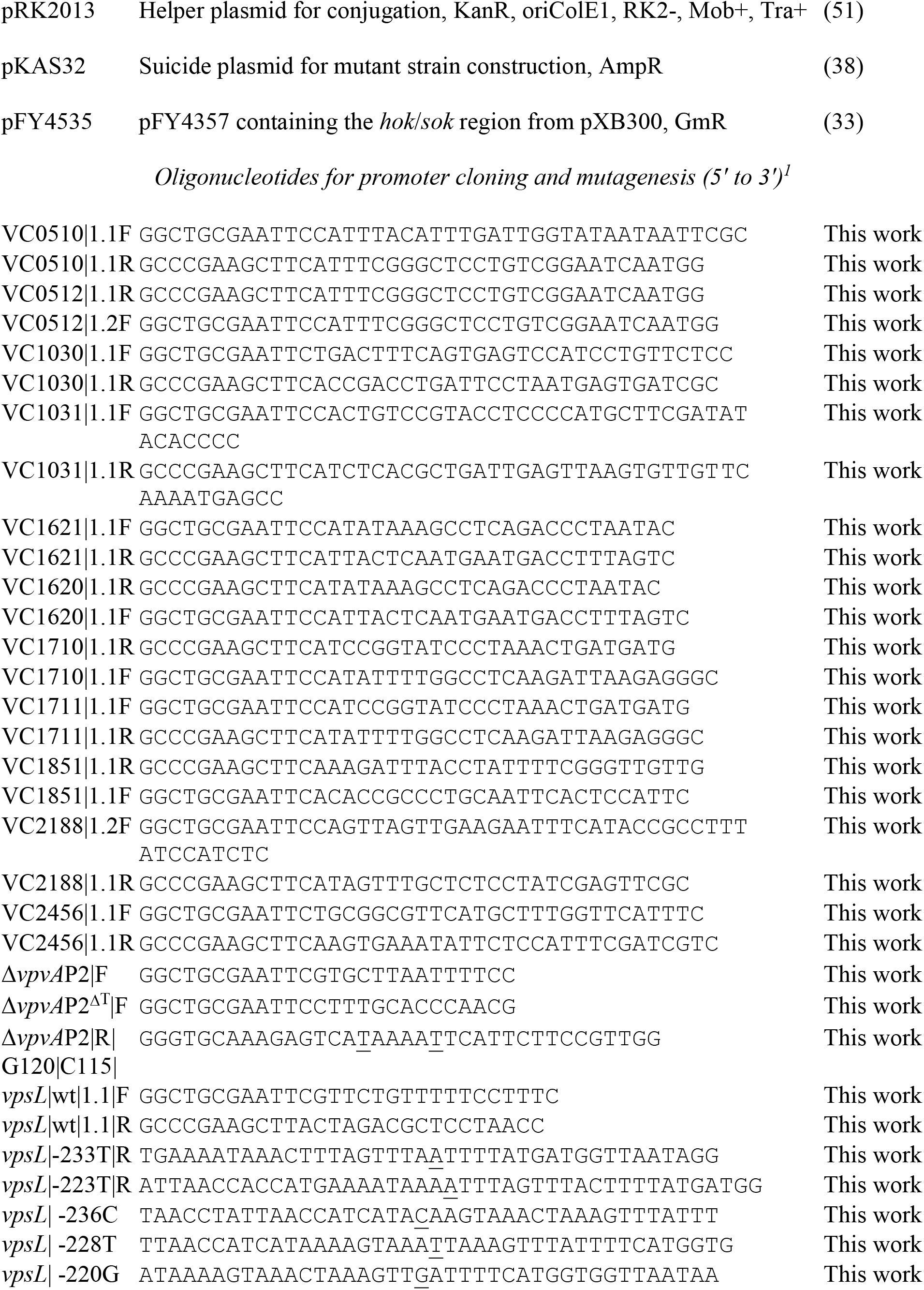
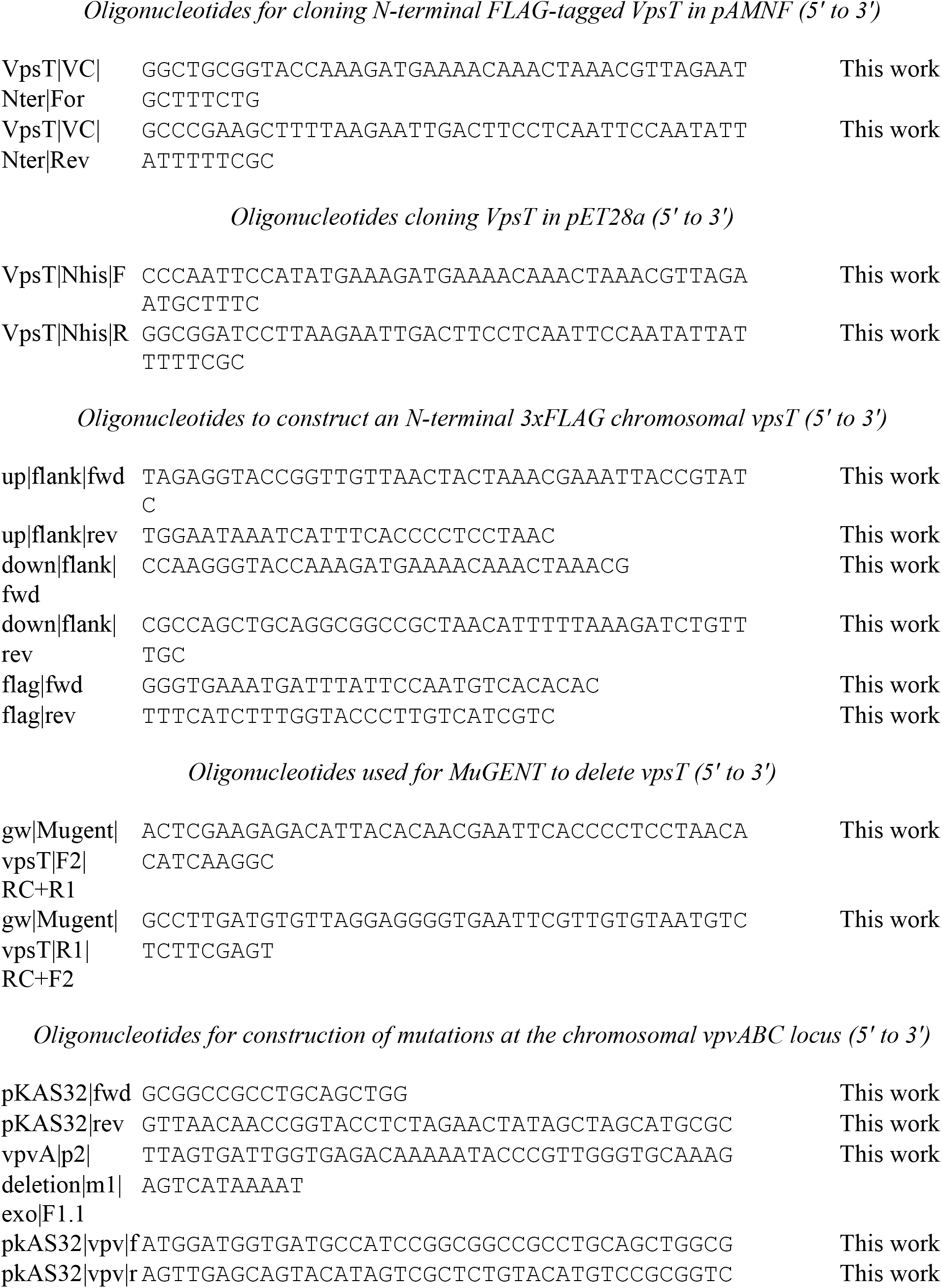
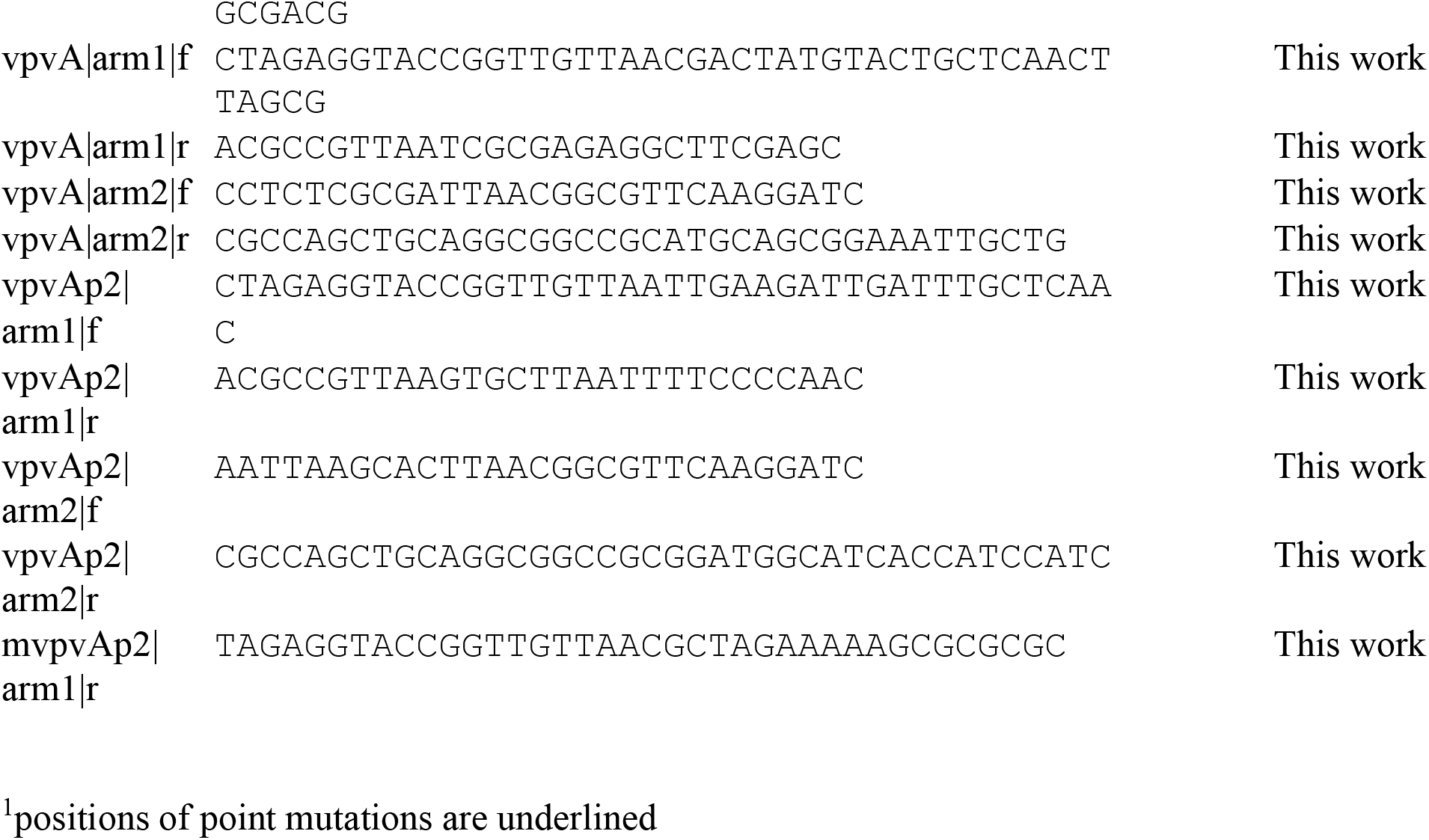
Strains, plasmids and oligonucleotides.

